# Mitochondrial protein interaction landscape of SS-31

**DOI:** 10.1101/739128

**Authors:** Juan D. Chavez, Xiaoting Tang, Matthew D. Campbell, Gustavo Reyes, Philip A. Kramer, Rudy Stuppard, Andrew Keller, David J. Marcinek, James E. Bruce

## Abstract

Mitochondrial dysfunction underlies the etiology of a broad spectrum of diseases including heart disease, cancer, neurodegenerative diseases, and the general aging process. Therapeutics that restore healthy mitochondrial function hold promise for treatment of these conditions. The synthetic tetrapeptide, elamipretide (SS-31), improves mitochondrial function, but mechanistic details of its pharmacological effects are unknown. Reportedly, SS-31 primarily interacts with the phospholipid cardiolipin in the inner mitochondrial membrane. Here we utilize chemical cross-linking with mass spectrometry to identify protein interactors of SS-31 in mitochondria. The SS-31-interacting proteins, all known cardiolipin binders, fall into two groups, those involved in ATP production through the oxidative phosphorylation pathway and those involved in 2-oxoglutarate metabolic processes. Residues cross-linked with SS-31 reveal binding regions that in many cases, are proximal to cardiolipin-protein interacting regions. These results offer the first glimpse of the protein interaction landscape of SS-31 and provide new mechanistic insight relevant to SS-31 mitochondrial therapy.

**Significance Statement:** SS-31 is a synthetic peptide that improves mitochondrial function and is currently undergoing clinical trials for treatments of heart failure, primary mitochondrial myopathy, and other mitochondrial diseases. SS-31 interacts with cardiolipin which is abundant in the inner mitochondrial membrane, but mechanistic details of its pharmacological effects are unknown. Here we apply a novel chemical cross-linking/mass spectrometry method to provide the first direct evidence for specific interactions between SS-31 and mitochondrial proteins. The identified SS-31 interactors are functional components in ATP production and 2-oxoglutarate metabolism and signaling, consistent with improved mitochondrial function resultant from SS-31 treatment. These results offer the first glimpse of the protein interaction landscape of SS-31 and provide new mechanistic insight relevant to SS-31 mitochondrial therapy.

## Introduction

Mitochondria are the power stations of the cell and generate cellular energy in the form of adenosine triphosphate (ATP). ATP is produced by oxidative phosphorylation (OXPHOS) through the mitochondrial electron transport system (ETS) comprising five multi-subunit protein complexes, complex I to complex IV, and complex V, ATP synthase. Mitochondrial dysfunction is associated with many human diseases which can manifest in any organ or tissue, including the nervous system, skeletal and cardiac muscles, kidneys, liver, and endocrine system, and present diverse clinical symptoms [1]. There are few effective treatments for mitochondrial disease as dysfunction of the ETS and OXPHOS for ATP production involves multiple proteins and multiple steps [2]. A conventional therapeutic strategy of targeting a single protein is unlikely to succeed in restoring function in complicated mitochondrial diseases. Another significant challenge for treating mitochondrial dysfunction is the efficient delivery of therapeutic molecules to the mitochondria. Elamipretide (SS-31) is a synthetic tetrapeptide that is a member of an emerging new class of therapeutics that selectively target mitochondria to restore mitochondrial bioenergetics. SS-31 is currently under clinical trials for multiple mitochondrial disorders including aging, mitochondrial genetic disease, ischemia, acute kidney injury, and heart failure [3–5]. SS-31 has shown promising results for treatment of various mitochondrial diseases, however, its mechanism of action remains unclear.

Mitochondria are composed of two membranes, an outer mitochondrial membrane (OMM) and an inner mitochondrial membrane (IMM) which forms invaginations, commonly named cristae. The cristae membranes constitute the backbone platform where mitochondrial respiratory complexes are located and OXPHOS takes place. Cardiolipin (CL), an unusual anionic lipid with two phosphate head groups and four acyl chains, is almost exclusively localized on the IMM [6]. It has long been known that CL has a critical role in bioenergetics, cristae morphology, and assembly of respiratory components into higher order supercomplexes [7, 8]. Bound CL molecules are required for the enzymatic activities and stabilities of both individual protein subunits and protein supercomplexes involved in mitochondrial respiration. For example, CL plays an essential role in the oligomerization of the c-rings and lubrication of its rotation in ATP synthase (CV), which can influence the stability of cristae structure through dimerization[9, 10]; CL acts as glue holding respiratory supercomplexes (CIII and CIV) together and steering their assembly and organization [11, 12]; the binding sites of CL identified close to the proton transfer pathway in CIII and CIV suggest a role of CL in proton uptake through the IMM [13–15]. In the case of the ADP/ATP translocase, ADT, three bound CL molecules securely anchor the carrier protein in the IMM and affect ADP/ATP transport activity [16]. Previous studies have shown that SS-31 peptide can penetrate the OMM and concentrate in the IMM by selectively binding to CL [3–5, 17]. The leading hypothesis is that the alternating aromatic-cationic motif of this synthetic tetrapeptide binds CL via dual interactions, i.e., hydrophobic interaction with acyl chain and electrostatic interactions with anionic phosphate head groups. This specific binding has been validated by fluorescence spectroscopy, isothermal titration calorimetry, and NMR analysis [18, 19]. Though not experimentally demonstrated, it was suggested that binding of SS-31 to CL helps induce tighter curvatures of cristae to stabilize the IMM structure and optimize the organization of respiratory chain supercomplexes for enhanced electron transfer and ATP production [5, 17]. Given the numerous lines of evidence showing SS-31-CL and CL-protein interactions, one could speculate direct interactions may exist between SS-31 and mitochondrial proteins. SS-31-protein interactions could play a direct or synergistic role with CL interactions in mitochondrial structure, cristae morphology, and ATP synthesis. To date, no empirical data exists to demonstrate SS-31 interacts with mitochondrial proteins, and the mechanistic details of how SS-31 modulates and rejuvenates mitochondrial function have remained elusive.

Chemical cross-linking mass spectrometry (XL-MS) has evolved to be a valuable and widely used tool for studying protein structures and interactions [20–23]. Recent development of protein interaction reporter (PIR) based XL-MS technology in our group has further enabled large-scale identification of protein interactions in complex mixtures from living cells [24–26] and isolated functional organelles [27, 28]. Previous studies have shown that SS-31, when added to permeabilized cells or isolated mitochondria, can decrease generation of H_2_O_2_, increase O_2_ consumption and ATP production [18, 19, 29–31]. To investigate the protein interaction landscape of SS-31, we utilized a N-terminal biotinylated form of SS-31 which allows for affinity enrichment of cross-linked SS-31-protein complexes. PIR-based cross-linkers were applied directly to isolated mitochondria incubated with biotinylated SS-31 (bSS-31) to secure a snapshot of protein interactions in their functional state. Subsequent mass spectrometry analysis uncovered the identities of SS-31 protein interactors with topological information. Here, we present the first report of the SS-31 protein interaction network, comprising twelve mitochondrial proteins belonging to nine enzymatic complexes. Interestingly, all twelve proteins were previously reported to bind with CL on the IMM and contribute directly and/or indirectly to mitochondrial respiration. These data provide the first evidence of direct interactions between SS-31 and CL-associated proteins. Thus, in addition to effects on IMM structure, the interaction between SS-31 and CL may serve to localize SS-31 to the IMM and facilitate its interactions with CL-binding proteins. These SS-31-protein interactions provide new insight into the functional role of SS-31 within mitochondria. Furthermore, our research establishes a novel and general approach to investigate the structural details of protein interactions with therapeutic molecules *in situ*.

## Results and Discussion

### bSS-31 interactome

SS-31 has been shown to selectively interact with CL, localizing and concentrating the peptide in the IMM [18]. To identify SS-31 protein interaction partners, we utilized an affinity tagged version containing a biotin group on the peptide N-terminus (bSS-31) (**Fig. S1A)** and carried out the experimental workflow illustrated in **Fig. 1**. Importantly, addition of a biotin tag did not affect SS-31 activity. SS-31 has been demonstrated to have no effect on mitochondrial function in young healthy mitochondria, while increasing mitochondrial ATP production and reducing redox stress in aged mouse muscle [30, 32]. Isolated mitochondria from aged mouse hearts were incubated with 10 μM bSS-31 for an hour prior to chemical cross-linking with a PIR cross-linker (DP-amide) **(Fig. 1, Fig. S1B**). After cross-linking, mitochondrial proteins were extracted with urea and subjected to a standard tryptic digestion protocol. Peptides cross-linked to bSS-31 were enriched using avidin affinity chromatography and subjected to LC-MS analysis employing two different acquisition methods, ReACT [33] and Mango [34] developed in our lab for the identification of PIR cross-linked peptide pairs. In total we were able to confidently identify 17 non-redundant cross-linked peptide pairs between bSS-31 and 16 Lys residues on 12 different proteins as shown in **Fig. 2 & Table S1**. The bSS-31 cross-linked peptide identifications were repeatedly observed across biological replicate samples originating from four mice. The cross-linked lysine residues are indicated in **Fig. 2**. Interestingly the bSS-31 interacting proteins can be grouped into two broad classes, those involved in ATP production and those utilizing 2-oxoglutarate (**Fig. 2**). Furthermore, each of the 12 proteins is known to directly interact with CL which is important in maintaining the structures and functions for these proteins (**Fig. 2**).

**Figure 1.**
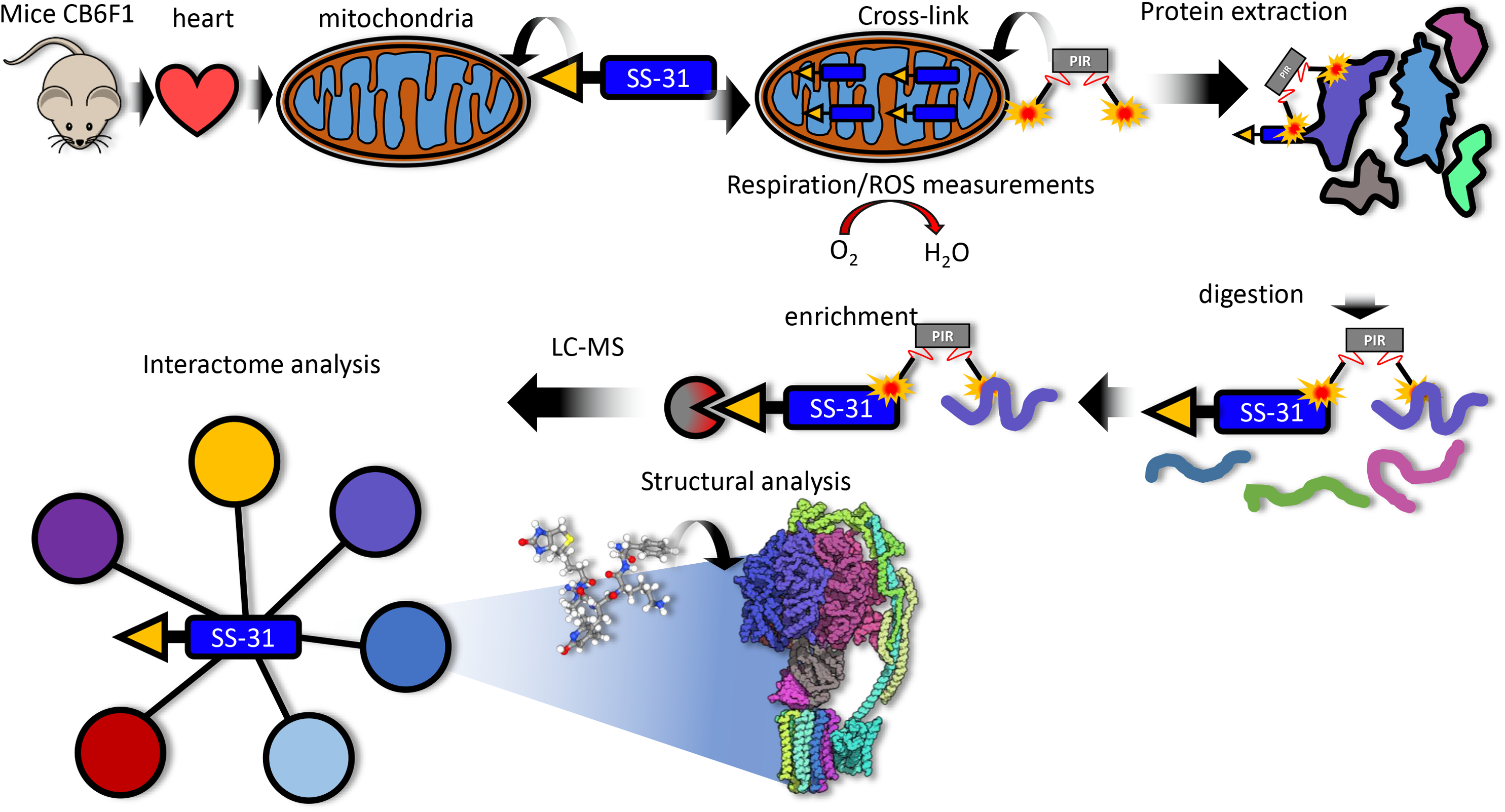
Experimental overview. Mice (total of 4) strain CB6F1 (BALB/cBy X C57BL/6) were euthanized by cervical dislocation. Hearts were excised and processed to isolate mitochondria. Isolated mitochondria were treated with 10 μM biotin SS-31 (bSS-31, blue rectangle with yellow triangle) for 1 h. Oxygen consumption rates and H_2_O_2_ production were monitored to evaluate the functional impact of bSS-31 (**Fig. S2**). Mitochondria were cross-linked with the PIR cross-linker DP-NHP (grey rectangle with orange stars). Protein was extracted using 8 M urea and digested with trypsin. Enrichment for peptides containing bSS-31 was performed with immobilized monomeric avidin. Peptide samples were analyzed by LC-MS to identify the interactome network for bSS-31 and structural analysis was performed using cross-link distance restraint guided molecular docking.

**Figure 2.**
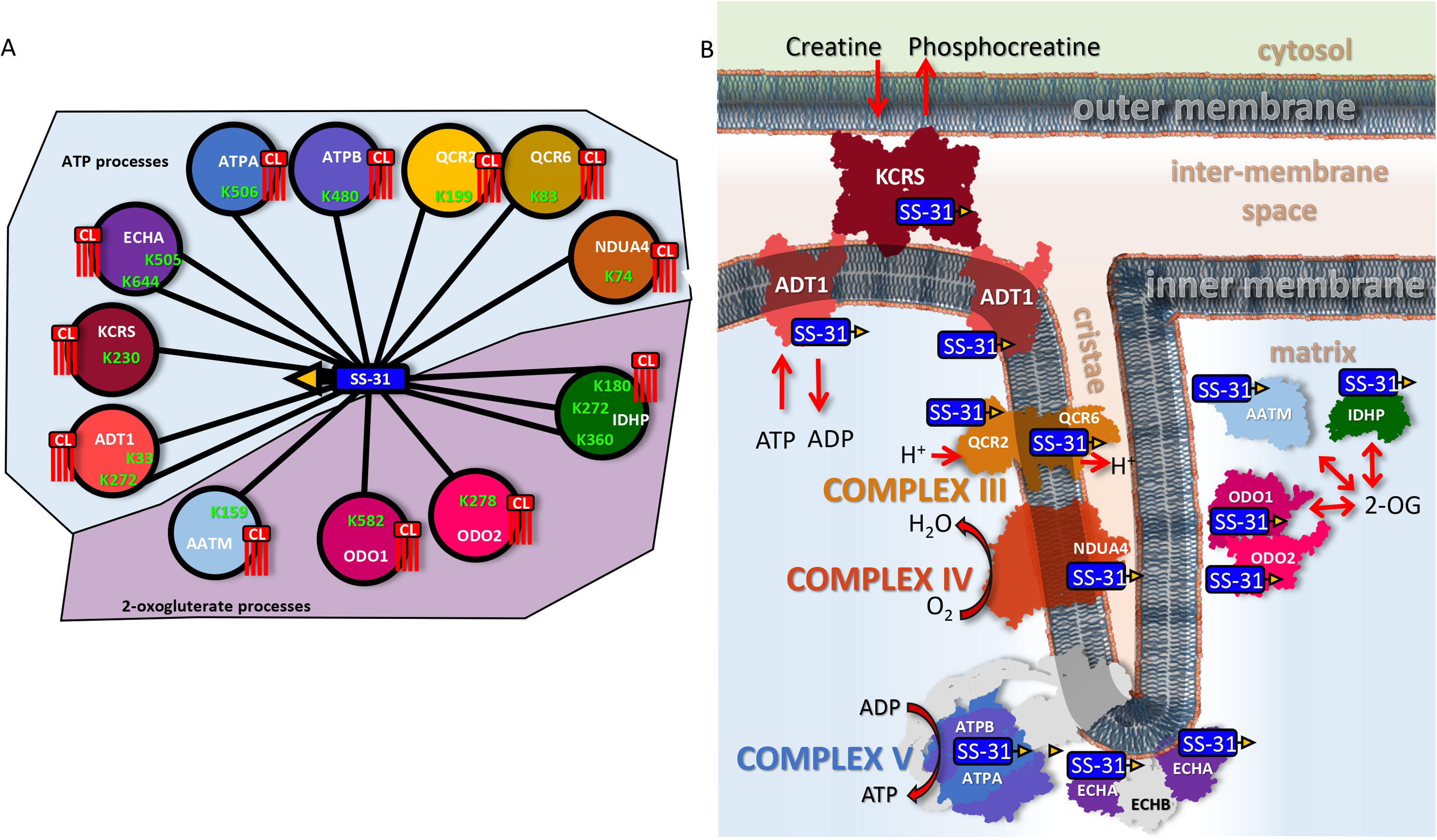
SS-31 interactome. **A)** Nodes represented by colored circles indicate the 12 proteins crosslinked to bSS-31. UniProt identifiers are labeled on the nodes along with the lysine residues that were identified as cross-linked to bSS-31. All 12 proteins are known to interact with cardiolipin (CL, red). The proteins are grouped into two major classes: ATP processes (upper light blue area) and 2-oxogluterate processes (lower light purple area). **B)** Schematic representation of SS-31 peptide cross-linked proteins within the mitochondria. These proteins include creatine kinase (KCRS), ADP/ATP translocase (ADT1), complex III (CIII) subunits QCR2 and QCR6, complex IV (CIV) subunit NDUA4, complex V (CV) subunits ATPA and ATPB, trifunctional enzyme subunit (ECHA), aspartate amino transferase (AATM), isocitrate dehydrogenase (IDHP) and 2-oxogluterate dehydrogenase complex subunits ODO1 and ODO2. bSS-31 peptide is represented by a blue rectangle with yellow triangle.

We utilized structural data from the Protein Data Bank (PDB) to localize the cross-linked residues and perform cross-link distance restraint guided molecular docking with bSS-31 using PatchDock [35]. For IDHP and AATM mouse structures were available in the PDB. For the other proteins we employed Clustal Omega [36] to align each mouse protein with homologous structures from other species. Note, all cross-linked Lys residues for the 10 mouse proteins for which no PDB structure exists are conserved in the homologous protein structures from other species. The data have been uploaded into XLinkDB [37] where a table of the cross-linked peptide pairs, interaction network and docked structures can be viewed (http://xlinkdb.gs.washington.edu/xlinkdb/BiotinylatedSS31_Bruce). For each protein docked with bSS-31, the interaction interface with bSS-31 was defined as those amino acid residues within 5 Å of the surface volume encapsulating the atom positions of bSS-31 from the top 10 scoring docked models. We utilized Composition Profiler [38] to compare the amino acid composition of the bSS-31 interfaces with that of the MitoCarta 2 (MC2) database [39]. Statistically-significant differences (P-value <0.05) included residues Asp and Thr which are enriched in bSS-31 interaction interfaces while Leu was depleted as compared with MC2 (**Fig. S2**). This observation suggests that electrostatic and hydrogen bonding interactions through Asp and Thr side chains are likely important for bSS-31-protein binding, consistent with the high number of cationic and H-bonding sites within bSS-31, as shown in cross-link-directed bSS-31-protein structural models discussed below.

### ETS complexes (CIII and CIV)

OXPHOS complexes generate and maintain the mitochondrial membrane potential (ΔΨ) through the ETS (CI-IV), which powers conversion of ADP to ATP by ATP synthase (CV). The activity of OXPHOS complexes is dependent on cristae morphology and interactions with CL for their proper assembly and function [7]. Excitingly, efforts presented here revealed bSS-31 cross-linked with multiple subunits of the OXPHOS complexes CIII, CIV and CV (**Fig. 2,3, and 4)** from aged mitochondria. Two subunits of CIII were cross-linked to bSS-31. These included K199 of QCR2 and K83 of QCR6 (PDB: 1sqp [40]). Interestingly, the two bSS-31 interacting subunits of CIII lie on opposite sides of the IMM with QCR2 on the electrochemically negative matrix side and QCR6 located on the electrochemically positive inter-membrane space side (**Fig. 3**). Structural details are presented in **Fig. 3**. CIII functions to reduce cytochrome c transferring electrons from coenzyme Q through the Q-cycle while shuttling protons across the IMM [41]. Structural analysis indicates CIII is normally assembled as a dimer (CIII_2_), either alone or as part of larger mitochondrial supercomplexes with varying copies of CI and CIV [42]. Each monomer of CIII is comprised of 11 subunits in higher eukaryotes [43]. QCR2 is a core subunit of CIII and is important for its dimerization. QCR6 is a highly acidic subunit that is important for the interaction of CIII with cytochrome c [44]. Reduced cytochrome c carries electrons from CIII to CIV. CIV is the terminal complex of the ETS ultimately transferring electrons from cytochrome c to reduce molecular oxygen to water. bSS-31 was cross-linked to K74 of the CIV subunit NDUA4 which is located in the inter-membrane space. Interestingly, the location of NDUA4 within CIV was only recently resolved as the 14^th^ subunit of the complex [45]. Its location within CIV, precludes CIV dimer formation, suggesting that in contrast to most crystal structures showing CIV assembled as a dimer of 13 subunit complexes, it is instead a 14 subunit monomer [45]. The 14 subunit monomer of CIV is also consistent with structures of the respirasome supercomplex (CI_1_CIII_2_CIV_1_) [45]. While not essential for assembly of other CIV subunits, NDUA4 plays a critical role in regulating CIV activity [46]. Monomeric CIV contains multiple CL binding sites that are important for enzymatic activity [47, 48]. Interestingly the PDB structure of human CIV with NDUA4 resolved recently (PDB: 5z62) indicates a CL molecule bound to a deep pocket formed between NDUA4 and cytochrome c oxidative subunits COX2 and COX3, suggesting it may be important for stabilizing the relatively weak interaction of NDUA4 with the rest of CIV [45].

**Figure 3.**
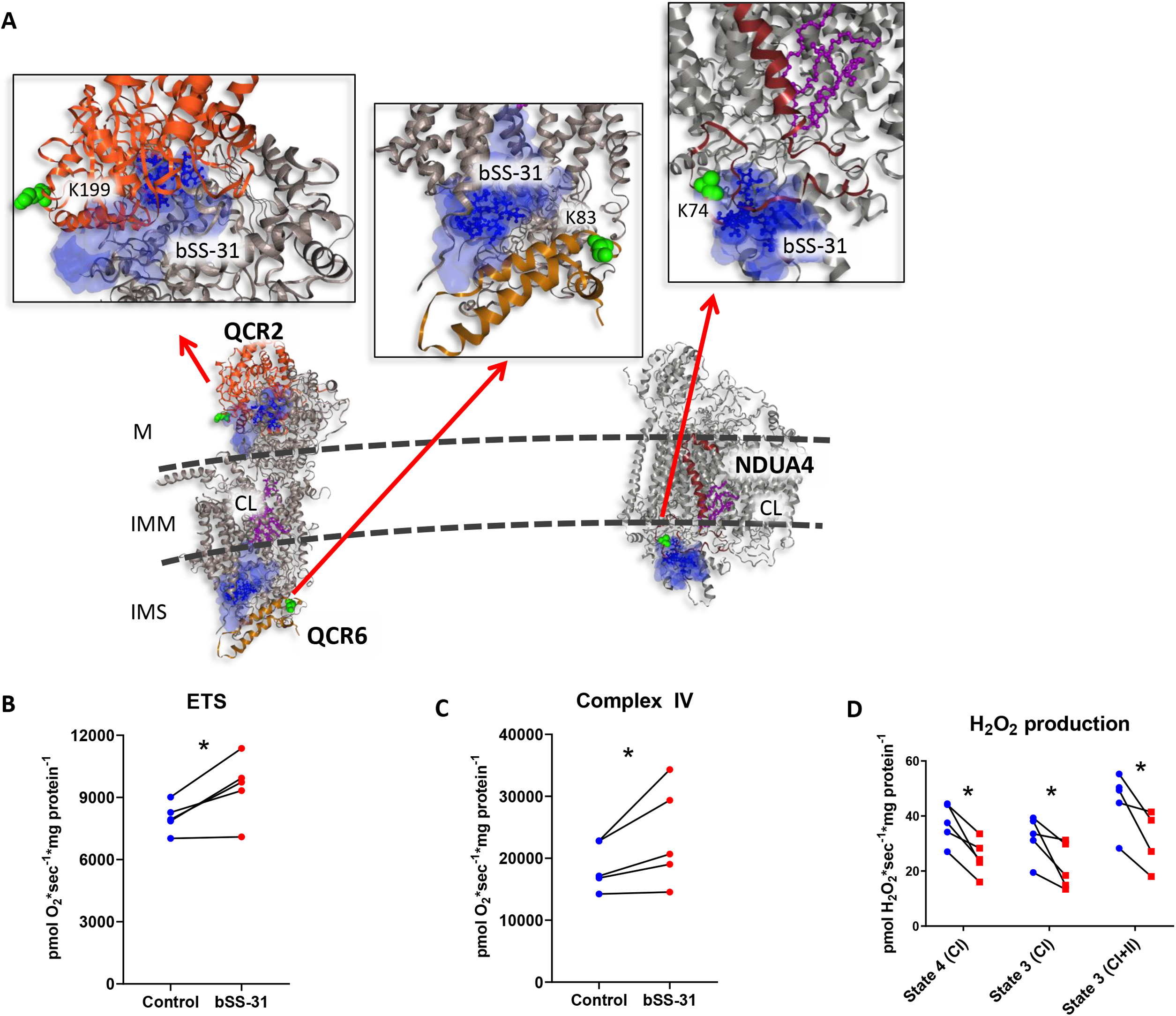
Structural and functional impact of bSS-31 interaction with OXPHOS complexes CIII and CIV. **A)** Structural view of interaction with subunits QCR2 and QCR6 of CIII and NDUA4 of CIV with inset showing zoomed view of docked structures. Distance constraints of 0-35 Å between the α-carbon of K3 of bSS-31 and the α-carbon of cross-linked lysine were used in molecular docking. Proteins are shown in cartoon view. Top ten molecular docking results for bSS-31 were included as shown in superimposed semi-transparent blue surface view with the top one dock position of bSS-31 shown in blue ball-and-stick representation. CL and CL binding residues are shown in magenta ball-and-stick and space fill representation, respectively. Cross-linked lysines are shown in green spheres. **B)** Maximum uncoupled respiration in mitochondria isolated from old mouse hearts in the presence of bSS-31 (red) or vehicle control (blue). **C)** Complex IV activity in the presence of bSS-31 (red) or vehicle control (blue). **D)** H_2_O_2_ production per O_2_ consumption in mitochondria isolated in the presence of bSS-31 (red) or vehicle control (blue). *P<0.05 using a paired t-test.

Functional measurements indicate that maximal flux through the electron transport system (ETS) in the presence of FCCP was significantly increased in mitochondria from old hearts incubated with bSS-31 compared to those without bSS-31 (**Fig. 3B)**, while there was no effect of bSS-31 in the mitochondria from young hearts (data not shown). The uncoupling by FCCP in the ETS measurement removes the dependence of respiration on CV. However, the ETS still provides an integrated measure of the electron transport system. In order to focus on one of the specific complexes that interact with bSS-31 we isolated CIV activity. Mitochondrial isolation performed with bSS-31 significantly elevated CIV specific activity **(Fig. 3C)** supporting a functional effect of the bSS-31 on this binding partner in mitochondria from aged hearts. While the enhanced ETS flux could be due to improved function of the ETS or increased substrate supply, the reduced H_2_O_2_ production reported here in **Fig. 3D** provided evidence for more efficient electron transfer through the ETS, i.e. fewer electrons escape before the reduction of O_2_ to water at CIV.

### ATP production and transport (CV, CK and ADT)

ATP synthase (CV), creatine kinase (CK), and ATP/ADP translocase (ADT) that we detected to crosslink with bSS-31 are three key components in participation of ATP production and transport. CV, ADT, together with inorganic phosphate carrier (PiC), form a larger supercomplex structure referred to as ATP the synthasome (**Fig. 2**) [49]. The ETS generated proton gradient and ΔΨ power the phosphorylation of ADP to ATP by CV. ADT and CK, two non-ETS proteins, play critical roles for shuttling metabolites to CV in the form of phosphocreatine (PCr) and ATP. The generation of ATP by OXPHOS is coupled to phosphorylation of creatine (Cr) by CK that utilizes ATP, transported via ADT protein, to synthesize PCr, which provides a temporal and special energy buffer for cells [50].

CV contains three copies each of ATPA and ATPB assembled as a heterohexamer in the F_1_ region of the complex on the matrix side of the IMM. There are six nucleotide binding sites located at the ATPA/ATPB interface. Cross-links were detected between bSS-31 and K506 of ATP synthase α-subunit (ATPA) and K480 of ATP synthase β-subunit (ATPB). Cross-link aided molecular docking localized bSS-31 to ATPB facing the IMM near the ATPA/ATPB interface (**Fig. 4A**). The CV dimer is thought to be essential to the formation of cristae by bending the IMM [51]. ATP synthase function depends on CL, which binds specifically, yet intermittently, to the IMM embedded c-ring portion of CV in the F0 region, acting as a lubricant for rotation of the CV rotor [10, 52, 53]. CV localizes near the tightly curved ends of cristae, where it assembles as rows of dimers stabilizing the IMM curvature [54–56]. Interestingly, isolated CV reconstituted into liposomes self-assemble into rows of dimers inducing membrane curvature [51]. Compartments generated by the membrane folds of cristae allow for pools of highly concentrated protons to drive CV rotation. Beyond the relevance of the dimer structure for ATP production by CV, CV dimers have also been implicated in formation of the mitochondrial permeability transition pore (mPTP) [57]. SS-31 has been shown to improve mitochondrial ATP production as well as inhibit mPTP formation [19, 31]. Prolonged treatment with SS-31 improved function in aged mitochondria, by increasing ATP production, improving flux of the ETS and reducing oxidative stress [32]. The protein interactors in CIII, CIV and CV identified here are central to all of these processes and may help explain the beneficial effects observed with SS-31 treatment.

**Figure 4.**
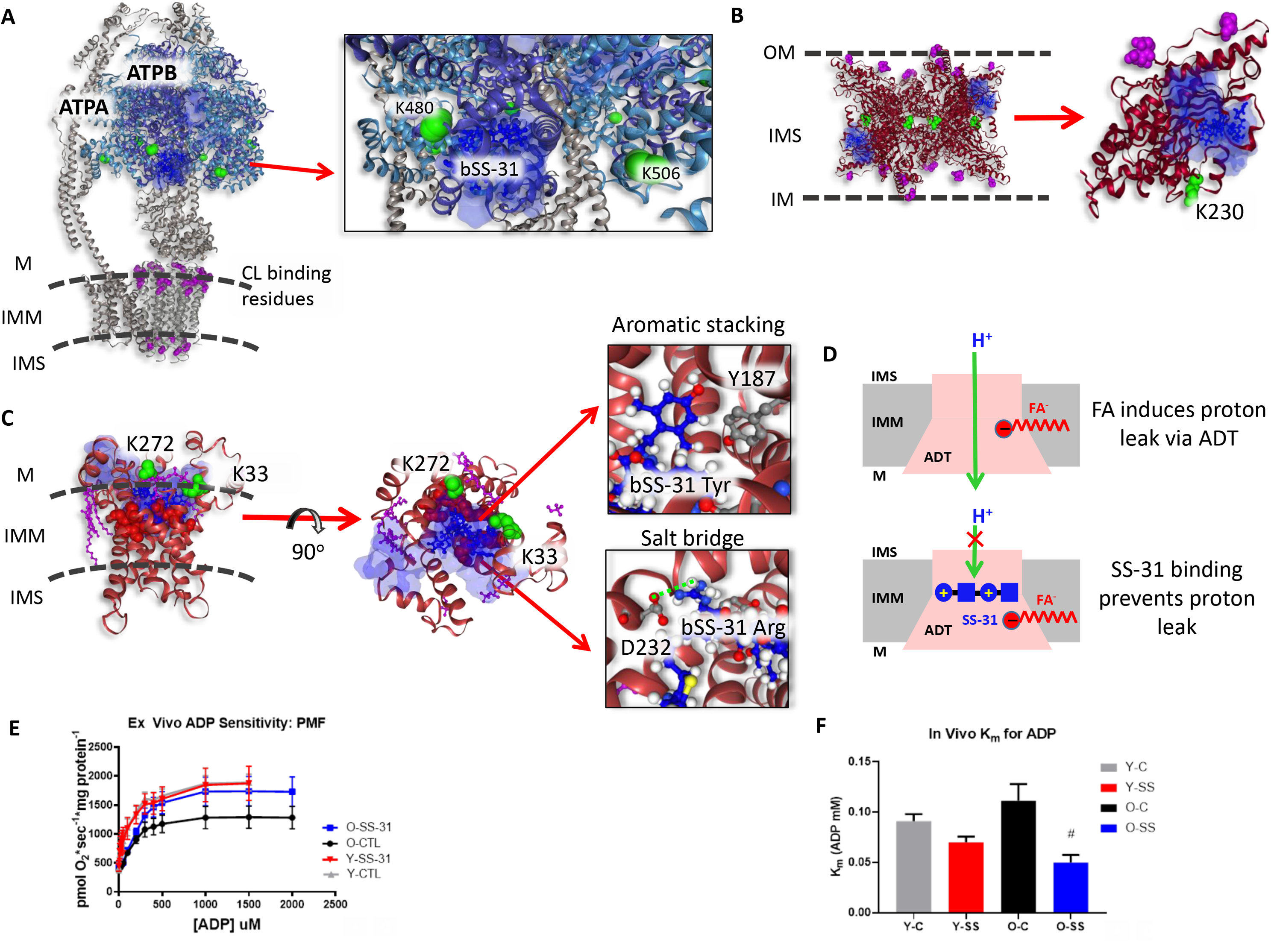
Structural and functional impact of bSS-31 interactions with ATP producing enzymes. Distance constraints of 0-35 Å between the α-carbon of K3 of bSS-31 and the α-carbon of cross-linked lysine residues (green spheres) were used in molecular docking. The top 10 docking results are selected to show the ranges of bSS-31 positions in semi-transparent blue surface view with the first docking position displayed in blue ball-and-stick view. CL and CL binding residues are shown in magenta ball-and- stick and space fill representation, respectively. **A)** Structure of bSS-31 interaction with ATP synthase with zoomed inset **B)** A side view of octameric KCRS (PDB: 1crk) bridging outer and inner membrane of mitochondria is shown in cartoon representation in left. Shown in right is monomeric chain A in ribbon view with CL binding residues, K408, R418, and K419, in magenta space-filled balls, and cross-linked residue K230 in green spheres. **C)** Structural representation of b-SS31 and ADT interaction in the BKA-locked m-state (PDB: 6gci). Cross-linked lysines K33 and K272 are shown in green spheres and CL is shown in magenta ball-and-stick. The substrate binding sites, K23, R80, R280, Y187, G183, I184, and S228, are shown in red spheres. Interactive versions of the structures can be viewed at http://xlinkdb.gs.washington.edu/xlinkdb/BiotinylatedSS31_Bruce. Insets show detailed view of salt bridge interaction between D232 residue and Arg residue in bSS-31 and aromatic stacking between Y187 and dimethyl tyrosine in bSS-31. **D)** Proposed model of FA-dependent proton leak via ADT shown on top [65], SS-31 binding of ADT possibly prevents proton leak with its two positively charged residues (bottom). **E)** SS-31 treatment increases ADP stimulated respiration in mitochondria isolated from old mice. **F)** SS-31 treatment decreases Km for ADP in vivo in old mice.

Creatine kinase S-type (KCRS) is a highly symmetric octamer localized in both intermembrane space and cristae space [58]. CL plays an important role for the binding of KCRS to the membranes. It has been suggested that this binding is stabilized by electrostatic interactions between the anionic phosphate head group in CL and a conserved basic motif, consisting of three conserved residues, K408, R418, and K419, in the C-terminal tail of KCRS [59]. The KCRS crystal structure (PDB: 1crk [60]) revealed two identical, opposite flat surfaces of the oligomers, allowing CK binding to two flat bilayer membranes between OMM and IMM, or the two segments of the cristae junctions (**Fig. 2 and 4B**) [60, 61]. The ability of KCRS to bridge two membranes via CL binding is an important factor for maintaining structural stability and integrity of mitochondria. This bridging is also important for the direct use of ATP, exported via the translocase, to phosphorylate Cr. Additionally, KCRS plays a critical role in clustering and enriching CL in the IMM and facilitating co-localization of ADP/ATP translocase into IMM. Our XL-MS results included the identification of bSS-31 interaction with KCRS at K230 (**Fig. 4B**). Superimposition of top ten molecular docking results localized bSS-31 peptide in the inner membrane space proximal to CL binding regions as displayed in blue cloud in **Fig. 4B**, consistent with KCRS binding CL and CL interaction with SS-31.

ADT, a transmembrane protein located in the IMM, is a carrier protein for importing ADP from the cytosol and exporting ATP from the mitochondrial matrix through cycling between cytoplasmic-open and matrix-open states (c-state and m-state) [62]. While the inhibitor carboxyatractyloside (CATR) locked ADT c-state crystal structure (PDB: 2c3e) [63] has been available for over a decade, better understanding of the mechanism of ADT for the full transport cycle became possible only recently with resolved crystal structure of bongkrekic acid (BKA) inhibitor locked ADT in m-state (PDB: 6gci) [64]. The six tilted transmembrane α-helices form a bucket-shaped cavity which opens to the intermembrane space (c-state) or the matrix space (m-state). Several CL molecules were found in the crystal structures, attached surrounding the bucket (**Fig 4C)**. CL molecules tightly bind through several hydrogen bonds and electrostatic interactions to act as an inter-domain bridge stabilizing ADT in the membranes [64]. The adenine nucleotide binding sites are found in the water-filled central cavity with the phosphate groups bound to K23, R80, R280; the adenine ring bound with Y187 via aromatic stacking; hydrophobic interactions with G183 and I184, and hydrogen bonding with S228 [62]. Our PIR cross-linking experiments identified ADT1, an isoform abundantly expressed in the heart, with two lysine residues (K33 and K272) cross-linked with bSS-31. The top ten molecular docking models produced using distance constraints from both lysine residues placed bSS-31 (shown in blue cloud) within the water-filled cavity of the translocase in m-state conformation (PDB: 6cgi) (**Fig. 4C**). bSS-31 is more likely to bind ADT in m-states for the following reasons. First, lysine residue K272 is only accessible for cross-linking in the m-state as it is exposed to the matrix side of the protein in m-states but buried in the membranes in c-states. Second, both lysine residues K33 and K272 are involved in salt bridge interactions in the c-state with D232 and E265, respectively, and are therefore less likely to be cross-linker reactive in the c-state. Moreover, in the top cross-link-directed docked bSS-31-ADT1 m-state model, D232 forms a salt bridge with the bSS-31 Arginine residue (**Fig. 4C**), and thus, would be unavailable for salt bridge formation with K33 that is important for c-state stabilization [64]. And finally, this model also locates the dimethyl tyrosine residue in bSS-31 proximal to Y187 and thus, aromatic stacking analogous to adenine ring interactions may stabilize bSS-31 in this m-state model as well (**Fig. 4C**).

Importantly, ADT was recently identified to support two distinct and competing transport modes involving ADP/ATP exchange and proton leak and thus, ADT connects coupled and uncoupled energy conversion mechanisms in mitochondria, respectively [65]. Studies have shown that proton leak through ADT was elevated in aged mitochondria and impairment of ADT was believed to play a central mechanism causing aged-related mitochondria defects [66–68]. More significantly, a recent report showed that mitochondrial trifunctional enzyme-deficient cardiac model system that simulates SIDS disease had a higher level proton leak and SS-31 treatment rescued the increased proton leak [69]; another recent report showed that SS-31 can suppress proton leak and rejuvenate mitochondrial function through direct binding with ADT [70]. SS-31 shares many common properties with inhibitors BKA and CATR such as larger molecular volume than ADP or ATP, and binding with ADT via multiple polar interactions [64]; and additionally, all three molecules were reported to block proton leak in ADT [70]. However unlike SS-31, BKA and CATR decreased the maximal respiratory rate and failed to enhance respiratory control ratio [70]. SS-31 binding to the ADT channel yet not functioning as an ADT inhibitor opens the possibility of therapeutic alteration of the balance of conformational states, potentially modulating proton leak and ADP/ATP exchange. As proposed in the recent report, the mechanism of proton leak through ADT is dependent on electrostatic interactions of the positively charged substrate binding site with a fatty acid (FA) co-factor, and competes with nucleotide transport through ADT [65]. Likewise, one can hypothesize that proton leak is prevented possibly through charge repulsion via two positively charged residues (Arginine and Lysine) in SS-31 (**Fig. 4D**). Since favorable SS-31 pharmacological effects on mitochondrial function include increased ATP production and decreased proton leak [3] [70], bSS-31 interaction with ADT in mitochondria identified in this work may be an important mediator of these beneficial effects. To assess the effect of SS-31 binding on the mitochondrial phosphorylation system we tested the effect of SS-31 on ADP stimulated respiration in mitochondria isolated from gastrocnemius muscles. State 3 respiration with complex I substrates was elevated in the presence of SS-31 in mitochondria from the aged gastrocnemius with no effect on mitochondria from the young **(Fig. 4E)**. Despite the increase in maximal respiration there was no effect of SS-31 on ADP sensitivity (Km) in isolated mitochondria. To further test the effect of SS-31 on ADP sensitivity in vivo we reanalyzed data from Siegel et al. [30] demonstrating that a single treatment with SS-31 *in vivo* increased ATPmax in aged mouse muscle **(Fig S3)**. The *in vivo* Km for ADP was significantly decreased in the aged mouse muscles following acute treatment with SS-31 with no effect in the young mice **(Fig. 4F**). This improved ADP sensitivity supports a functional effect of SS-31 on either ADT1, CV or both.

In addition to direct binding of SS-31 with ADT, the same group also reported that SS-31 associated with ATP synthasome and stabilized it [70]. Our crosslink data provides direct evidences that bSS-31 binds with two members of ATP synthasome supercomplex, CV and ADT. Since favorable SS-31 pharmacological effects on mitochondrial function include increased ATP production and decreased proton leak [3] [70], bSS-31 interaction with CV, ADT, and CK involved in mitochondria respiration identified in this work may be an important mediator of these beneficial effects.

### Trifunctional enzyme

The mitochondrial trifunctional enzyme is a hetero-oligomeric complex localized on the IMM that catalyzes the latter three reactions of fatty acid β-oxidation for very long chain fatty acids. The trifunctional enzyme consists of the α-subunit (ECHA) which contains 2-enoyl-CoA hydratase (ECH), and 3-hydroxyacyl-CoA dehydrogenase (HAD) activities, and the β-subunit (ECHB) containing 3-ketothiolase (KT) activity. Within mitochondria, the trifunctional enzyme is thought to primarily be assembled as an α2β2 hetero-tetramer. Recently, structural models for the human trifunctional enzyme have been determined by cryo-EM [71] and x-ray crystallography [72] both highlighting important structural differences between the mammalian mitochondrial trifunctional enzyme and the structures of bacterial orthologs [73, 74]. Both structures for the α2β2 mitochondrial trifunctional enzyme reveal a curved structure to the complex where the concave surface is important for interaction with the IMM [71, 72]. Beyond functioning in fatty acid β-oxidation, ECHA also functions to remodel CL via monolysocardiolipin acyltransferase activity [75]. The α-subunit of the mitochondrial trifunctional enzyme (ECHA) was cross-linked at K505 and K644 to bSS-31. Both of these residues exist on the concave side of the complex near the inner mitochondrial membrane where cardiolipin resides (PDB: 6dv2) (**Fig. 5**). Cross-linking distance constraint guided docking of bSS-31 with ECHA places bSS-31 into a substrate channel in ECHA near the HAD active side, proposed to be important for cardiolipin remodeling [72] (**Fig. 5**). Interestingly SS-31 was recently shown to rescue excessive proton leak in cardiomyocytes with mutated ECHA [69].

**Figure 5.**
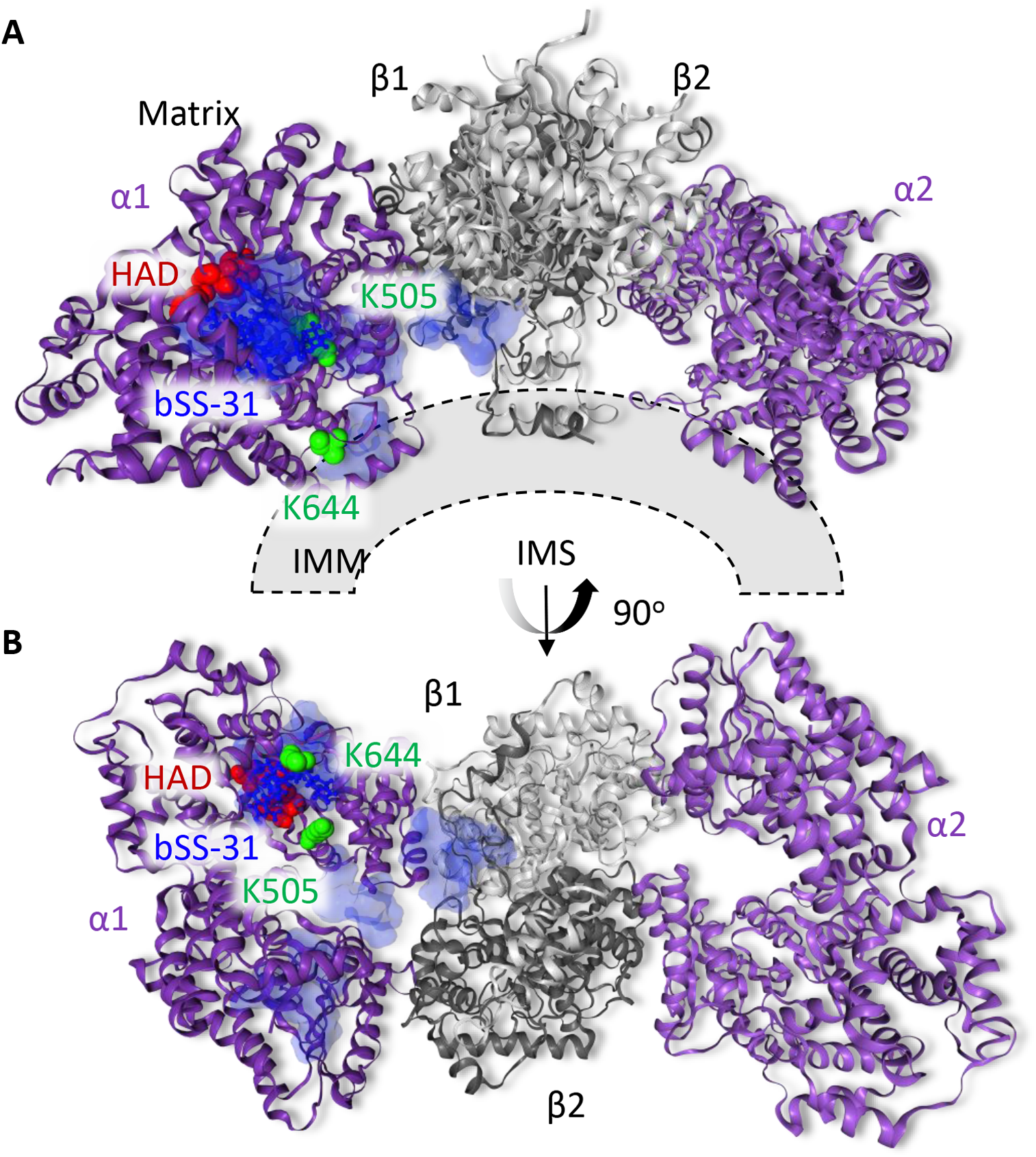
Structural view of bSS-31 interaction with ECHA. Distance constraints of 0-35 Å between the α-carbon of K3 of bSS-31 and the α-carbon of cross-linked lysine residues K644 and K505 of PDB structure 6dv2 were used in molecular docking. The top 10 docking results are selected to show the ranges of bSS-31 positions in semi-transparent blue surface view with the first docking position displayed in blue ball- and-stick view. The cross-linked Lys residues on ECHA are shown as green space filled residues. The HAD active site residues proposed to be important for cardiolipin remodeling (S477, H498, E510, T548) are shown as red space filled residues. Interactive versions of the structures can be viewed at http://xlinkdb.gs.washington.edu/xlinkdb/BiotinylatedSS31_Bruce.

### 2-oxoglutarate enzymes

The metabolite 2-oxoglutarate (aka α-ketoglutarate, 2-OG) is a key intermediate of the tricarboxylic acid The 2-OGDHC is a large multi-protein assembly(TCA) cycle, building block for amino acids and nitrogen transporter. Due to its involvement in both carbon and nitrogen metabolism and multiple signaling pathways, 2-OG has been suggested to act as a master regulator metabolite [76]. One example of this is highlighted by the discovery that 2-OG can extend the lifespan of *Caenorhabditis elegans* by inhibiting ATP synthase and target of rapamycin (TOR) downstream [77]. 2-OG also plays a key role in hypoxia signaling serving as a substrate for prolylhydroxylase (PHD) enzymes which regulate the stability of hypoxia inducible factors (HIF) transcription factors [78, 79]. Interestingly, we identified bSS-31 cross-linked with multiple lysine sites across four key proteins including isocitrate dehydrogenase, 2-OG dehydrogenase (ODO1), dihydrolipoyllysine-residue succinyltransferase component of 2-OG dehydrogenase complex (ODO2) and aspartate aminotransferase (AATM), all of which produce or utilize 2-OG in their reactions. Excitingly these interactions suggest a potential role for SS-31 in 2-OG signaling.

Isocitrate dehydrogenase catalyzes the decarboxylation of isocitrate forming 2-OG and carbon dioxide. There are three types of isocitrate dehydrogenase in mammals, IDH1, IDH2 and IDH3, which differ in structure, function and subcellular localization [80]. IDH1 and IDH2 are NADP^+^-dependent enzymes that catalyze reversible reactions and are localized in the cytosol and mitochondria respectively [80]. IDH3 is the NAD^+^-dependent form, localized to the mitochondrial matrix functioning primarily in energy production in the TCA cycle where it catalyzes an irreversible reaction and is allosterically regulated by a number of cofactors [80]. The mitochondrial localized NADP^+^-dependent IDH2 (aka IDHP) also functions in the TCA cycle but has additional roles in supplying NADPH used in redox homeostasis and fatty acid metabolism [81]. This places IDHP at a critical crossroad, regulating energy production through the TCA cycle, maintaining the mitochondrial pool of NADPH needed for the glutathione antioxidant system and producing 2-OG utilized by PHD proteins in HIF1α signaling. Interestingly, IDHP can also interact with CL containing membranes that can induce a conformational change in IDHP regulating its enzymatic activity [82]. Cross-links were identified between bSS-31 and K180, K272 and K360 of IDHP. Docking of bSS31 with the PDB structure of IDHP (5h3f) results in localization of bSS-31 to a groove between the large and small domains of IDHP near the isocitrate binding pocket (**Fig. 6A&B**). Electrostatic repulsion between positively charged Lys residues located in the large domain (K127, K129, K130 and K133) and the small domain (K256) has been suggested to play an important role in IDHP activity by widening the cleft between the domains [83] (**Fig. 6B**). Neutralization of the charged residues by acetylation or mutation on these Lys residues results in a narrower cleft, smaller isocitrate binding pocket and lower enzymatic activity [83]. A number of acidic residues surround the bSS-31 interface, potentially participating in electrostatic interactions with bSS-31 (**Fig. 6B**). It is possible that when SS-31 interacts with IDHP, the positive charges in SS-31 would also play a role in widening the cleft between the domains, potentially impacting IDHP activity.

Catalyzing the rate limiting fourth step of the TCA cycle is 2-OG dehydrogenase complex (2-OGDHC), which utilizes thiamine pyrophosphate (TPP) as a cofactor and reduces NAD^+^ to NADH during the conversion of 2-OG to succinyl-CoA and CO_2_. The 2-OGDHC is a large multi-protein assembly (~2 MDa) localized on the IMM comprising multiple copies of three enzymatic components E1 (ODO1), E2 (ODO2) and E3 (dihydrolipoyl dehydrogenase, DLDH) and sharing similar assembly with the pyruvate dehydrogenase complex (PDH) and the branched chain 2-oxoacid dehydrogenase complex (BCKDH). There are no high-resolution structural models available for any components of the mammalian 2-OGDHC. Experimental evidence from multiple sources including genetic mutational, limited proteolysis and electron microscopy indicate the 2-OGDHC is composed of 24 E2 subunits assembled into a cubic core with varying numbers of E1 and E3 subunits bound to the periphery of the core [84]. The activity of the 2-OGDHC is tightly regulated, being inhibited by ATP, NADH and succinyl-CoA, intertwining its function with that of the OXPHOS complexes. Under some conditions it can also be a primary source of mitochondrial ROS production producing superoxide and hydrogen peroxide at much higher rates than complex I, which is often implicated as the primary source of mitochondrial ROS [85]. The E3 component of 2-OGDHC produces ROS in the form of H_2_O_2_ and superoxide radicals when electrons from the FADH_2_ transfer to molecular oxygen. Furthermore, very reactive and damaging lipoate thiyl radicals can form in the E2 component as a result of reaction with a FADH semiquinone [85]. We identified cross-links between bSS-31 and K582 of ODO1 (E1) and K278 of ODO2 (E2) **Fig. 6C**. Interestingly, CL has been shown to be important for the stability of the 2-OGDHC [86].

**Figure 6.**
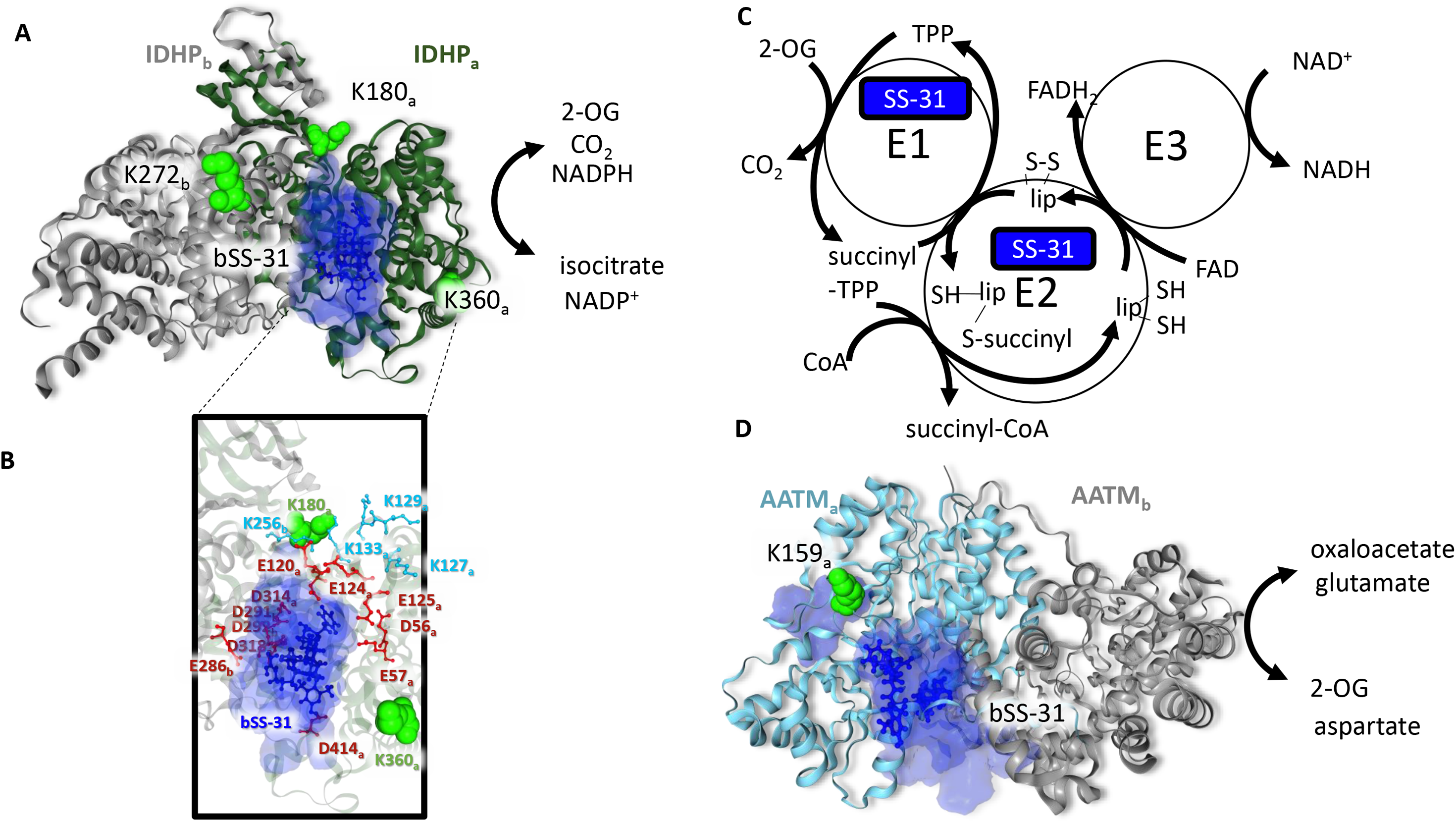
SS-31 interaction with 2-oxoglutarate enzymes. **A)** bSS-31 interaction with IDHP (PDB 5h3f) indicated as a homodimer with the IDHPa chain shown as a green ribbon and the IDHPb chain in grey. Lysine residues cross-linked to bSS-31 are displayed as green space filled residues and are labeled with subscript “a” or “b” to indicate which IDHP monomer they are on. bSS-31 (blue ball and stick structure) was docked onto the a-subunit using distance restraints for K180 and K360 from the a-subunit and K272 from the b-subunit. The volume encompassing the positions of bSS-31 from the top 10 docking results is displayed as a semi-transparent blue surface. **B)** View of the bSS-31 interaction interface highlighting surrounding acidic residues (red) and Lys residues (cyan) which are important in maintaining the groove between the large and small domains of IDHP [83]. **C)** Schematic representation of the 2-oxoglutarate dehydrogenase complex (2-OGDHC). bSS-31 was identified as cross-linked to ODO1 a component of the E1 of 2-OGDHC and ODO2 a component of the E2 portion of 2-OGDHC. **D)** bSS-31 interaction with AATM (PDB 3pdb) shown as a homodimer with the a-chain in teal and the b-chain in grey. Lysine K159 (green space filled residue) of the a-chain was used to dock bSS-31 into the structure. The volume encompassing the positions of bSS-31 from the top 10 docking results is displayed as a semi-transparent blue surface. Interactive versions of the structures can be viewed at http://xlinkdb.gs.washington.edu/xlinkdb/BiotinylatedSS31_Bruce.

The mitochondrial aspartate aminotransferase (AATM) catalyzes the reversible reaction of aspartate and 2-OG to glutamate and oxaloacetate. AATM is a key enzyme linking amino acid and carbohydrate metabolism by providing a carbon source (2-OG) to the TCA cycle as well as nitrogen for the synthesis of non-essential amino acids and nucleotides [87]. Beyond its canonical function, mitochondrial AATM also functions as kynurenine aminotransferase as well as a transporter for long chain fatty acids [87, 88]. AATM also interacts directly with the cardiolipin containing liposomes, the inner-mitochondrial membrane as well as the 2-OGDHC [89, 90]. We identified a cross-link between bSS-31 and K159 of AATM. Cross-link guided docking places bSS-31 near the oxaloacetate binding pocket in the PDB structure 3PDB **Fig 6D**. This is on the opposite side of AATM from the LCFA binding pocket, colored as blue space filled residues in **Fig 6D**.

Overall the enzymes IDHP, ODO1, ODO2 and AATM function in closely connected biological pathways related to regulation of 2-OG levels, mitochondrial redox homeostasis and HIF1-α signaling in addition to key metabolic pathways. The specific interactions detected between these proteins and bSS-31 suggest a possible role for SS-31 to influence this nexus of mitochondrial function.

Previous efforts to elucidate SS-31 mechanisms have suggested that beneficial therapeutic effects arise from SS-31 interaction with CL. While CL binding may be the primary impact of SS-31 treatment, the present work resulted in identification of the first set of SS-31 interacting proteins from intact, functional mitochondria using PIR XL-MS. These results provide new insight and new opportunities to better understand the molecular actions of this mitochondrial targeted therapeutic. This set of 12 proteins are all part of a larger energy transfer system for mitochondrial ATP production through the OXPHOS pathway. Four of the proteins are directly involved with 2-OG metabolism and signaling which regulates mitochondrial metabolism, redox homeostasis, and cellular response to hypoxia. Importantly each of the SS-31 interacting proteins also interact with CL, which impacts their structure and function. This observation presents the possibility that SS-31 concentrates in the IMM by interacting with CL, facilitating its interactions with components of the OXPHOS and 2-OG signaling pathways. Numerous studies have demonstrated the beneficial effects of SS-31 to restore healthy mitochondrial function, increase ATP production, and reduce mitochondrial redox stress. Beyond identifying protein interaction partners, the XL-MS data provide structural information, in terms of distance restraints, that were used to localize bSS-31 interacting regions on these proteins. Knowledge of these protein interactors lays the foundation for follow up functional studies to evaluate the significance of these interactions. As a general method, employing affinity tagged drugs with XL-MS should be extensible to investigate the interaction landscapes of therapeutic molecules in complex biological samples such as organelles, cells and tissues.

## Materials and Methods

### Animal husbandry

This study was reviewed and approved by the University of Washington Institutional Animal Care and Use Committee. Male, 36-37 month old CB6F1 (BL/6: BalbC) mice were received from the National Institute of Aging aged mouse colony. All mice were maintained at 21°C on a 14/10 light/dark cycle and given standard mouse chow and water ad libitum with no deviation prior to or after experimental procedures. Animals were sacrificed by cervical dislocation with no anesthetic.

### Extraction of tissue, mitochondrial isolation and SS-31 treatment

On day of animal sacrifice, the heart (120-150 mg wet weight) was resected. The whole heart was homogenized on ice in a glass Dounce homogenizer in 2 mL of ice-cold respiration buffer (RB, 1 mM EGTA, 5 mM MgCl2, 105 mM K-MES, 30 mM KCl, 10 mM KH_2_PO_4_, pH 7.1). The homogenates were centrifuged for 10 min at 800 × g at 4 °C. The supernatants were collected and centrifuged for 15 min at 8,000–10,000 × g at 4 °C. After the removal of supernatant, the pellet was resuspended in isolation medium. The pellet was centrifuged for 15 min at 8,000–10,000 × g at 4 °C and was resuspended in isolation medium. Mitochondrial pellets were separated into new centrifuge tubes and diluted with 200 μL RB, 10 mM Pyruvate, and 2 mM Malate. Either elamipretide or 2 μL saline was added to the mitochondria and incubated for 1 hour at room temperature with mild rocking.

### Ex Vivo Mitochondrial Respiration and H_2_O_2_ Production

In a parallel experiment, mitochondria were isolated from young (5-7 month-old) and old (36-37 month old) mouse hearts or gastrocnemius muscles as described above either in the presence or absence of 10 μM bSS-31. Approximately 100 μg mitochondrial homogenate was used in the 2 mL chamber of an Oxygraph 2K dual respirometer/fluorometer (Oroboros Instruments, Innsbruck, Austria) at 37°C and stirred gently during substrate and inhibitor titrations. Heart mitochondrial respiration and H_2_O_2_ production were measured simultaneously under the following conditions. State 4 was measured by adding 5 mM pyruvate, 2 mM malate, and 10 mM glutamate. State 3 was stimulated by adding 2.5 mM ADP (CI) followed by 10 mM succinate (CI+CII) then cytochrome C (6 mM). The rate of maximal uncoupled flux through the ETS was measured by adding 1 μM FCCP. Following FCCP, Antimycin A (2.5 uM) was added to block flux through complex III followed by TMPD and ascorbate (1mM and 4mM, respectively) to measure complex IV activity. The non-mitochondrial rate of oxygen consumption was subtracted from all measured functional parameters before reporting final values. In a separate experiment ADP sensitivity was measured in gastrocnemius muscles by measuring complex I stimulated respiration across a range of ADP conenctrations.H_2_O_2_ emission was measured in parallel with respiration using Amplex Red (10 mM) and HRP (0.1 U/mL).

### In vivo ADP sensitivity assays

In order to assess in vivo ADP sensitivity we reanalyzed data from Siegel et al. [30]. Briefly, a short ischemic period was used to induce PCr breakdown. This rate of PCr breakdown is equal to the resting mitochondrial ATP production. The PCr recovery was measured over 6 min to determine a time constant of recovery (t_PCr_) to yield ATPmax(= PCr_rest_/ t_PCr_). Free ADP concentration at rest and the start of recovery and ATP production at the start of recovery (ATP_flux_= ΔPCr/ t_PCr_) used to determine ADP sensitivity by fitting the [ADP] and ATPase rates at rest and initial recovery to Michaelis-Menton plots with a hill coefficient of 2.6 (**Fig. S3**)[30, 91].

### Cross-linker synthesis

The protein interaction reporter (PIR) cross-linker amide-DP-NHP was synthesized by solid phase peptide synthesis using a CEM Liberty Lite peptide synthesizer. Amino acids were coupled to amide-Rink resin in the following order: biotin-Lys, Fmoc-Lys(Fmoc), Fmoc-Pro, Fmoc-Asp, succinic anhydride. The N-hydroxyphthalamide (NHP) ester of trifluoroacetic acid (TFA) was synthesized by dissolving 5.86 g of NHP in 20 mL of TFA anhydride in a 50 mL round bottom flask. The reaction was allowed to proceed for 1.5 h under a dry N_2_ atmosphere with constant mixing via magnetic stir bar. After 1.5 h the mixture was dried under vacuum to obtain a white crystalline solid (TFA-NHP). The cross-linker (0.5 mmoles of peptide on resin) was esterified by reacting with a 12-fold molar excess of TFA-NHP in 10 mL of dry pyridine for 20 min at room temperature with constant mixing. The reaction mixture was transferred to a Bio-Rad poly prep column and the liquid was filtered away. The resin containing the esterified peptide was washed extensively with a total of 60 mL dimethyl formamide (DMF) followed by 60 mL of dichloromethane (DCM). The cross-linker was cleaved from the resin by incubation with 5 mL of 95% TFA, 5% DCM for 3 h at room temperature with constant mixing. The cross-linker was precipitated by adding the cleavage solution to ice cold diethyl ether at a ratio of 1:15 by volume. The cross-linker was pelleted by centrifugation at 3400 g for 30 min at 4°C. The cross-linker pellet was washed by resuspending the pellet in 10 mL of fresh ice cold diethyl ether and repeating the centrifugation step. The cross-linker pellet was then dried by vacuum centrifugation and weighed. The cross-linker was dissolved in DMSO to a concentration of 270 mM, aliquoted and stored at −80°C until used.

### Cross-linking reaction

The cross-linker amide-DP-NHP was added at a final concentration of 10 mM (from a 270 mM stock solution) to mitochondria incubated with bSS-31. The reaction mixture was mixed at 800 rpm at room temperature for 30 min. A total of 5 cross-linked samples were generated including 4 biological replicates of mouse heart mitochondria incubated with bSS-31 and a negative control sample to which no bSS-31 was added. After cross-linking the samples were frozen at −80°C.

### Cross-linked mitochondrial sample preparation

Cross-linked mitochondrial samples were lysed with 8 M urea in 0.1 M NH_4_HCO_3_. The sample was kept on ice and sonicated with a GE 130 ultrasonic processor using 5 cycles of 5 s pulses at an amplitude of 40. The total protein concentration was measured using the Pierce Coomassie Plus Bradfrod protein assay. Protein disulfide bonds were reduced with 5 mM TCEP for 30 min followed by alkylation of thiol groups with 10 mM iodoacetamide for 30 min. Samples were diluted 10-fold with 0.1 M NH_4_HCO_3_ and digested with a 1:200 ratio of trypsin to protein at 37°C overnight. After digestion samples were acidified by adding trifluoroacetic acid (TFA) to a final concentration of 1%. Acidified samples were desalted using Waters C18 Sep Pak cartridges and associated vacuum manifold. Samples were passed through the cartridge at a flow rate of 1 mL/min, followed by 3 mL washes of water containing 0.1% TFA. Peptides were eluted from the Sep Pak cartridge with 1 mL of 80% acetonitrile, 20% water, 0.1% TFA. Desalted samples were concentrated by vacuum centrifugation using a Genevac EZ-2 system. Dried samples were reconstituted in 0.5 mL of 0.1 M NH_4_HCO_3_ and adjusted to pH 8 with 1.5 M NaOH. 100 uL of 50% UltraLink monomeric avidin slurry was added to the sample and incubated with constant mixing for 30 min at room temperature. The monomeric avidin resin was washed 5 times with 3 mL of 0.1 M NH_4_HCO_3_ followed by elution of bound peptides with 2 additions of 0.5 mL of 70% acetonitrile, 0.5% formic acid, each with 5 min incubation. The eluate fractions were pooled and concentrated by vacuum centrifugation. The biotin enriched sample was reconstituted in 30 uL of 0.1% formic acid in water and subjected to LC-MS analysis as described below.

### LC-MS analysis of cross-linked peptide pairs

Samples were analyzed by LC-MS using two methods developed for analysis of cross-linked peptide pairs, namely ReACT [33] and Mango [34]. ReACT analysis was carried out on a Velos-FTICR mass spectrometer coupled with a Waters nanoAcquity UPLC. Peptides were fractionated by reversed-phase liquid chromatography by first loading the sample (5 μL injection) onto a trapping column (3 cm x 100 um i.d.) packed with Reprosil C8 beads (Dr. Maisch) using a flow rate of 2 μL/min of 98% solvent A (water, 0.1% formic acid) and 2% solvent B (acetonitrile, 0.1% formic acid). After trapping peptides were separated over an analytical column (60 cm x 75 um) maintained at 45°C, packed with Reprosil C8 stationary phase using a flow rate of 300 nL/min and applying a binary gradient of 90% solvent A / 2% solvent B to 60% solvent A / 40% solvent B over 120 min followed by a wash cycle consisting of 20% solvent A / 80% solvent B for 20 min and a re-equilibration period consisting of 98% solvent A / 2% solvent B for 20 min. Eluting peptides were ionized by electrospray ionization (ESI) by applying a voltage of 2.6 kV to a laser pulled spray tip at the end of the chromatography column. MS1 analysis (500-2000 m/z) was performed in the ICR cell with a resolving power setting of 50K at 400 m/z, and an automatic gain control (AGC) setting of 5E5. The most abundant precursor ion with a charge state ≥ 4 was selected for MS2 using an isolation window of 3 m/z and a normalized collision energy (NCE) of 25. Fragment ions were analyzed in the ICR cell using a resolving power setting of 12.5K at 400 m/z and an AGC setting of 2E5. MS2 spectra were searched in real-time for fragment ions that satisfy the expected PIR mass relationship (mass peptide 1 + mass peptide 2 + mass reporter = mass precursor) within a 20 ppm mass error tolerance. If satisfied the two released peptide ions were sequentially analyzed by MS3 in the Velos dual ion trap mass analyzer where they were isolated with a 3.0 m/z isolation window and fragmented with a NCE of 35 using an AGC setting of 5E4.

Samples were analyzed by Mango using a Thermo Q-exactive plus mass spectrometer coupled with a Thermo Easy nLC. 5 μL of sample was injected into the nLC system where it was loaded onto a trapping column (3 cm x 100 um i.d.) packed with Reprosil C8 beads (Dr. Maisch) using a flow rate of 2 μL/min of solvent A. After trapping peptides were separated over an analytical column (60 cm x 75 um) maintained at 45°C, packed with Reprosil C8 stationary phase using a flow rate of 300 nL/min by applying the same binary gradient described for the ReACT analysis above. Eluting peptides were ionized by electrospray ionization (ESI) by applying a voltage of 2.6 kV to a laser pulled spray tip at the end of the chromatography column. MS1 analysis (400-2000 m/z) was performed in the orbitrap mass analyzer using a resolving power setting of 70K at 200 m/z and an ACG value of 1E6. This was followed by MS2 on the 5 most abundant precursor ions with charge states ≥ 4 using a 3 m/z isolation window, a resolving power setting of 70K at 200 m/z and an ACG value of 5E4.

### Data analysis

LC-MS data files in. RAW format were converted to .mzXML format using the ReADW tool in the Trans Proteomic Pipeline software suite [92]. Comet [93] was used to search the .mzXML files against the Mitocarta 2 database [39] containing both forward and reverse protein sequences (2084 total sequences) along with the addition of the sequence for bSS-31 (Biotin-D-Arg-dimethyl Tyr-Lys-Phe-NH2). Due to the presence of non-canonical amino acids the bSS-31 sequence was entered using the following single letter amino acid sequence BJKZ. Within the Comet parameters file the mass of B (Biotin-D-Arg) was set to 382.17926 Da, the mass of J (dimethyl Tyr) was set to 191.094629 Da and the mass of Z (amidated Phe) was set to 146.084399 Da. Additional Comet parameters used for searching ReACT data included; a peptide mass tolerance of 20 ppm, allowance of −1/0/1/2/3 ^13^C offsets, trypsin as the digesting enzyme considering only fully tryptic sequences with up to 5 missed cleavage sites. Oxidation of Met (15.9949 Da) was included as a variable modification while the cross-linker stump mass modification on Lys (197.032422 Da) was included as a required modification at any position within the peptide sequence except for the C-terminus, MS3 spectra were searched using a 1.0005 Da tolerance on the fragment ions with a bin offset of 0.4. Comet parameters used for Mango data were the same except that the Mango search parameter was set to 1, MS2 spectra were searched and a 0.02 Da tolerance with a 0.0 bin offset were used for fragment ions.

### Validation of bSS-31 fragmentation spectrum

Ten μL of 10.6 mM bSS-31 in 18.2 MΩ H_2_O was transferred to a 1.5 mL tube. 90 μL of 170 mM Na_2_HPO_4_ pH 8.0 was added. This was followed by addition of 1.11 μL of 270 mM DP-amide cross-linker stock solution in DMSO resulting in a 3 mM final cross-linker concentration. The reaction was carried out at room temperature for 1 h with shaking 1400 rpm. The sample was then acidified to 1 % TFA by volume. The sample was then desalted using a 50 mg size C18 Sep-Pak column. After loading the sample onto the Sep-Pak the peptides were washed with 3 additions of 1 mL H_2_O/0.1% TFA. The sample was then eluted from the Sep-Pak column with 1 mL 50% ACN/ 2% CH3COOH and analyzed by direct infusion MS with the Velos-FTICR mass spectrometer. SpectraST v 5.0 was used to search fragmentation spectra generated by ReACT against a spectral library of the cross-linker modified bSS31. Fragmentation spectra assigned as bSS-31 cross-links were required to contain the accurate mass of the DP stump modified bSS-31 (Fig. S1C) and have a SpectraST assigned p-value of less than 0.1.

### Molecular docking

A molecular model for bSS-31 was generated using Avogadro (v. 1.2.0) utilizing a molecular mechanics conformer search of 1000 conformers followed by geometric optimization. PatchDock [35] was used to dock the model of bSS-31 with structural models in the PDB for bSS-31 interacting proteins (AATM = PDB: 3pdb, ADT c-state = PDB:2c3e, ADT m-state = PDB: 6gci, ATPA/ATPB = PDB: 5ARA, QCR2/QCR6 = PDB: 1sqp, NDUA4 = PDB: 5z62, ECHA = PDB: 6dv2, IDHP = PDB: 5h3f, KCRS = PDB: 1crk). Clustal Omega [36] was used to align each mouse protein with homologous structures from other species. For each protein, the top 10 scoring docked models from PatchDock were downloaded and the interaction interface with bSS-31 was defined as those amino acid residues within 5 Å of the surface volume encapsulating the atom positions of bSS-31.

## Supplemental Figure Legends

**Fig. S1.**
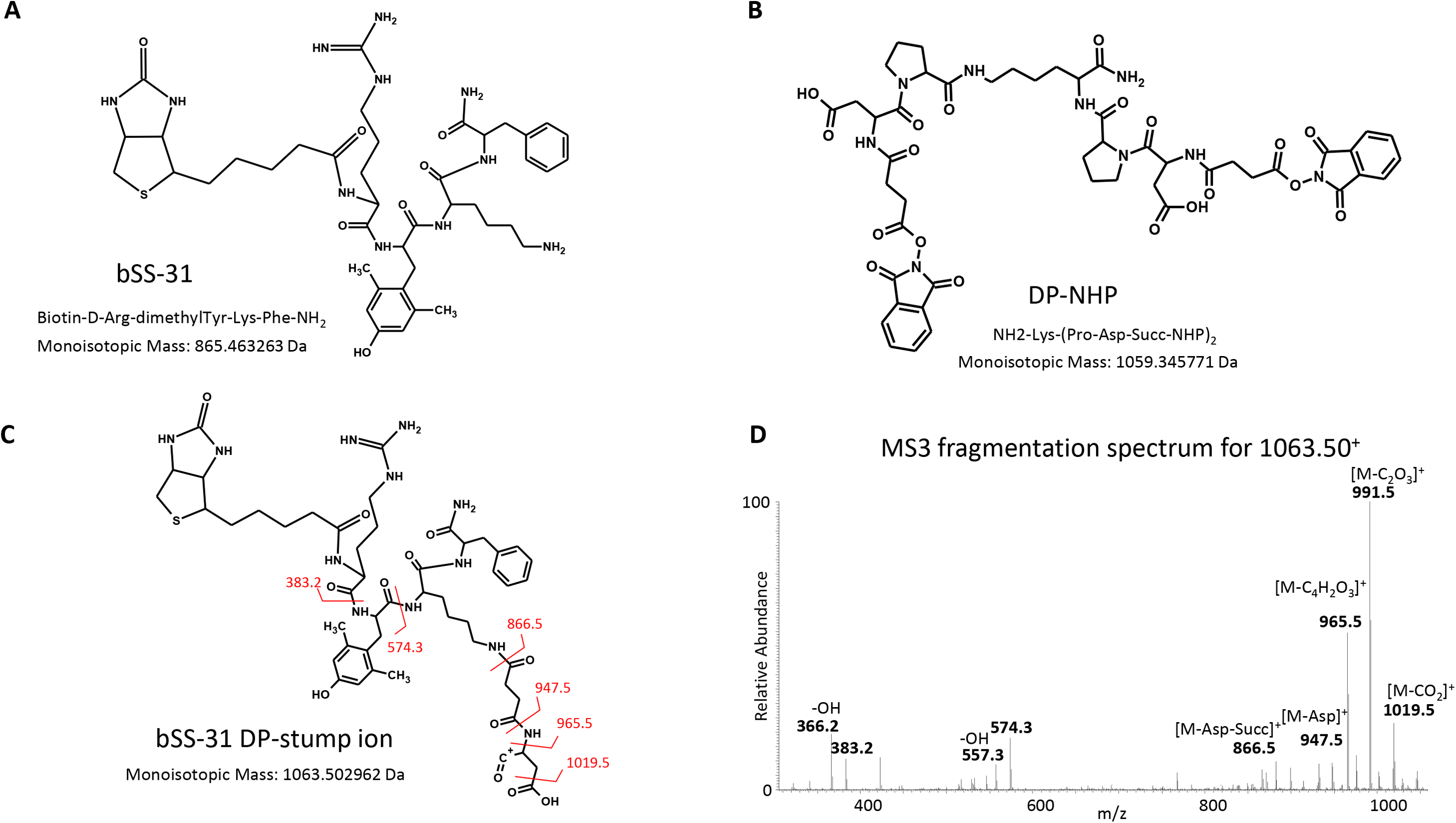
Structures of biotin SS-31 and PIR cross-linker DP-NHP. **A)** chemical structure of biotinylated SS-31. **B)** chemical structure of DP-NHP cross-linker. **C)** chemical structure for bSS-31 DP-stump ion which results from MS2 fragmentation of the PIR labile bonds of DP cross-linked bSS-31. Red lines with numbers indicate the bods which break to give rise to the major fragment ions observed in panel D. **D)** MS3 fragmentation pattern for bSS-31 DP stump ion with major fragment ions labeled. Bonds which fragment to give rise to these fragment ions are indicated in red in panel C. Ions at m/z 557.3 and m/z 366.2 result from loss of hydroxyl radical from fragments ions at m/z 574.3 and m/z 383.2 respectively.

**Fig. S2.**
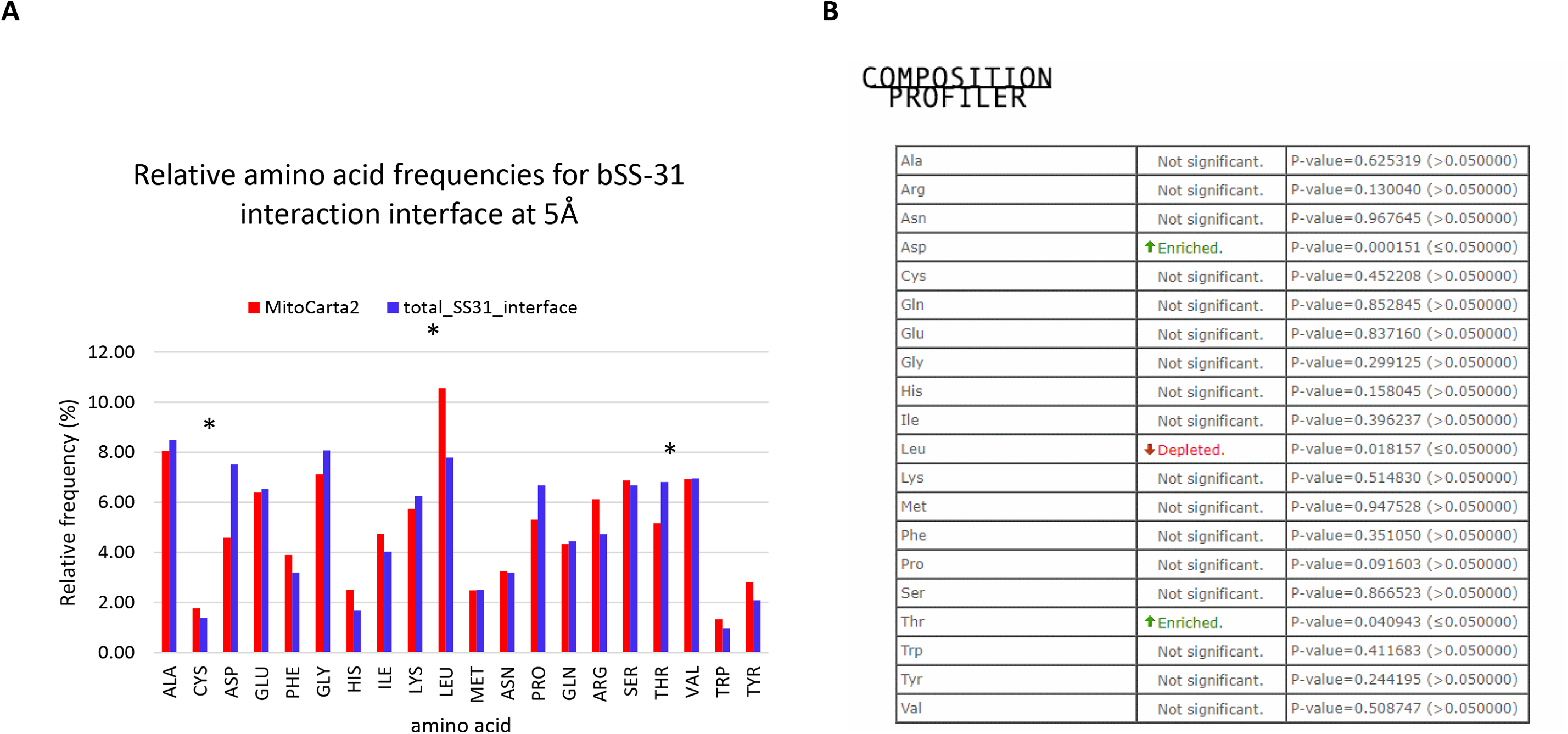
Amino acid composition of bSS-31 interaction interfaces – **A)** Bar plot indicating relative frequencies for the 20 amino acids in the MitoCarta2 database (red) and the amino acids comprising the bSS-31 interaction interface (residues within 5 angstroms of bSS-31 in docked models). Asterisks indicate a statistically significant difference (p<0.05) as calculated by the Composition Profiler tool. **B)** Output from Composition Profile comparing amino acid composition of the bSS-31 interaction interfaces to the MitoCarta2 database.

**Fig. S3.**
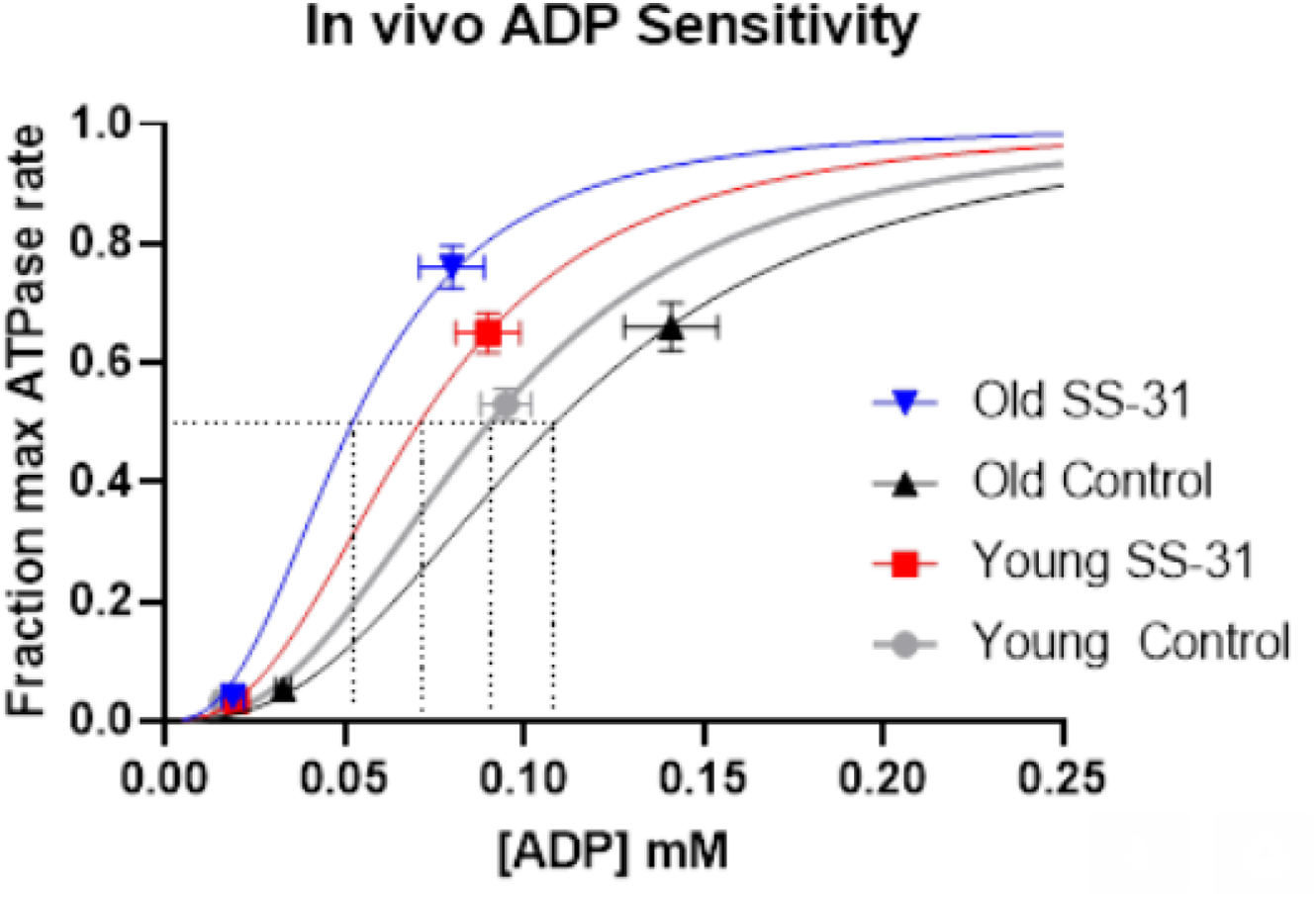
Effect of SS-31 on ADP sensitivity in vivo. Free ADP concentration at rest and the start of recovery and ATP production at the start of recovery (ATPflux= ΔPCr/ tPCr) used to determine ADP sensitivity by fitting the [ADP] and ATPase rates at rest and initial recovery to Michaelis-Menton plots with a hill coefficient of 2.6

**Table S1 –** List of identified bSS-31 cross-linked peptides

**Video S1 –** Cross-link directed docked models for bSS-31 ADT1 interaction in the c-state (PDB: 2c3e) and m-state (PDB: 6gci). ADT is shown as red ribbon structures with the cross-linked residues K33 and K272 displayed as yellow space filled residues. Electrostatic interactions between D232-K33 and E265-K272 stabilize the c-state of ADT and are not present in the m-state. In the m-state docking results, the top scoring model without atomic clashes, suggests a potential charge interaction between D232 and the Arg of bSS-31.

## Acknowledgments

The authors would like to thank Hazel Szeto for her generous gift of biotin labeled SS-31. We would also like to thank the members of the Bruce Lab, Peter Rabinovitch and Nathan Alder for helpful input during these experiments. This work was supported by P01 AG001751, an American Federation for Aging Research BIG award, 5R01GM086688, 5R01RR023334, 1R01GM097112, 7S10RR025107, 5U19AI107775, and 1R01HL110879.

